# Effects of stimulus timing on the acquisition of an olfactory working memory task in head-fixed mice

**DOI:** 10.1101/2022.08.26.502106

**Authors:** Josefine Reuschenbach, Janine K Reinert, Izumi Fukunaga

## Abstract

Knowing what factors affect the acquisition of a behavioural task is central to understanding the mechanisms of learning and memory. It also has practical implications, as animal behavioural experiments used to probe cognitive functions often require long training durations. Delayed Match (or Non-Match)-to-Sample (DMS/DNMS) tasks are relatively complex tasks used to study working memory and sensory perception, but their use in the mouse remains hampered by the lengthy training involved. In this study, we assessed two aspects of stimulus timing on the acquisition of an olfactory DNMS task: how the sample-test odour delay durations and the reward timing affect the acquisition rate. We demonstrate that head-fixed mice learn to perform an olfactory DNMS task more quickly when the initial training uses a shorter sample-test odour delay without detectable loss of generalisability. Unexpectedly, we observed a slower task acquisition when the odour-reward interval was shorter. This effect was accompanied by a shortening of reaction times and more frequent sporadic licking. Analysis of this result using a drift-diffusion model indicated that a primary consequence of early reward delivery is a lower decision bound. Since an accurate performance with a lower decision bound requires greater discriminability in the sensory representations, this may underlie the slower learning rate with early reward arrival. Together, our results reflect the possible effects of stimulus timing on stimulus encoding and its consequence on the acquisition of a complex task.

## Introduction

How animals learn is a question that has long intrigued humankind (Hume, 1779; Pavlov, 1927; Skinner, 1950; Thorndike, 1898). Systematic studies of behavioural changes resulting from stimulus associations have enabled quantitative descriptions of relationships between physical stimuli and behaviour and in some cases, allowed identification of corresponding physiological mechanisms (Milner et al., 1998). Insights from such animal experiments impact human interactions, too, as seen in their influence on educational psychology (Kay and Kibble, 2016; Shuell, 1986). In neuroscience research, there is a practical consequence, as animal experiments to probe cognitive functions often require lengthy training paradigms, which is especially true when the acquired behaviours are not innate.

DMS/DNMS tasks are relatively complex tasks in that the association formed is not simply between one stimulus with a reward (Blough, 1959; Skinner, 1950). They are paradigms where animals are presented with two stimuli (”sample” and “test” stimuli), separated by an interval (”delay”), and the task for the animals is to report if the identities of the two stimuli match in the case of DNM task or are different, in the case of DNMS task. These paradigms have become essential for studying the working memory in various model organisms, ranging from pigeons and dolphins to primates and humans (Blough, 1959; Hampson et al., 1993; Herman and Gordon, 1974; Mishkin and Delacour, 1975; Skinner, 1950). They have also been used to probe the perceptual similarities of sensory stimuli (Nakayama and Rinberg, 2021; Zentall and Smith, 2016) and continue to reveal insights into brain functions, such as the roles played by the sensory and prefrontal cortices (Eriksson et al., 2015; Liu et al., 2014; Wu et al., 2020; Zhang et al., 2019) and the limbic system (Eichenbaum et al., 2007; Hampson et al., 1993; Mishkin and Manning, 1978) in retaining information over time.

Despite the usefulness and the long history of these paradigms, their use in the mouse still faces a bottleneck due to the lengthy training required for accurate performance (Goto et al., 2010; Liu et al., 2014; Nakayama and Rinberg, 2021; Wu et al., 2020). For example, in the delayed match-to-position, which is a spatial working memory task in freely-moving mice, more than 25 sessions are required to reach high accuracy (Goto et al., 2010). Several recent studies succeeded in training head-fixed mice to perform olfactory DMS/DNMS tasks, though the training protocols described vary considerably in length and procedure. For example, in one study that trained head-fixed mice on a DMS paradigm, a median of 25 days’ training was reported to acquire the task and involved minor punishment to suppress unwanted, sporadic licking during the delay phase (Wu et al., 2020). Other studies included an auto-shaping phase, which involves using only stimuli associated with a reward, without other types of trials (Han et al., 2018; Liu et al., 2014; Nakayama and Rinberg, 2021). Some of these studies report a robust acquisition in 5 days following the initial auto-shaping stage (Han et al., 2018; Liu et al., 2014), though this required multiple training sessions per day (Han et al., 2018). There is also variation in the inter-odour (sample-test odour) interval used in the initial training, ranging from 1.5 seconds to 5 seconds (Han et al., 2018; Liu et al., 2014; Nakayama and Rinberg, 2021; Wu et al., 2020; Zhang et al., 2019). Thus, it is presently unclear what factors are essential for fast and robust acquisition of the DNMS in head-fixed mice and if a simpler design suffices.

In this study, we wished to understand the factors that lead to efficient DNMS task learning. In particular, we focused on two aspects of stimulus timing. First, we sought to assess the effect of the inter-odour interval on the rate of DNMS acquisition since short-term retention of information tends to decay with increasing delay intervals (Baddeley, 2011). Training with a shorter sample-test interval is expected to be more efficient, but without the guarantee that the performance generalises well to longer intervals. In addition, we assessed the timing of the reward, or the unconditioned stimulus, as some theoretical studies predict that longer stimulus-reward intervals lead to reduced reinforcement learning, for example, through a weakening of the “eligibility trace” over time (Barto et al., 1983).

We demonstrate that the acquisition of the DNMS task is quicker when the initial training uses a shorter inter-odour delay without detectable loss of generalisability. Unexpectedly, we observed a slower task acquisition with a shorter stimulus-reward interval. In combination with a drift-diffusion model, we reveal that a primary characteristic of early reward delivery during training is a lowering of the decision bound. This lower bound requires greater discriminability in the sensory representations to achieve the same behavioural accuracy, which may underlie the slower learning rate. Together, our results reflect on possible consequences of stimulus timings on the quality of stimulus encoding and the acquisition of a complex task.

## Results

In the DNMS task, two stimuli (”sample” and “test” stimuli) are presented with a delay in between, and the animals are required to report when the identities of the two stimuli do not match. To implement this task using olfactory stimuli, an olfactometer should be capable of presenting stable olfactory stimuli with minimal cross-contamination, irrespective of the sample-test delay durations used. To achieve this, we designed a novel olfactometer (Fig. 1; see Methods). Stable and clean presentations of the two odours are achieved by separating the air streams for the sample and test odours for most parts of the odour pulse preparation while allowing continuous air passages to avoid a pressure build-up (Fig. 1A).

**Figure 1:**
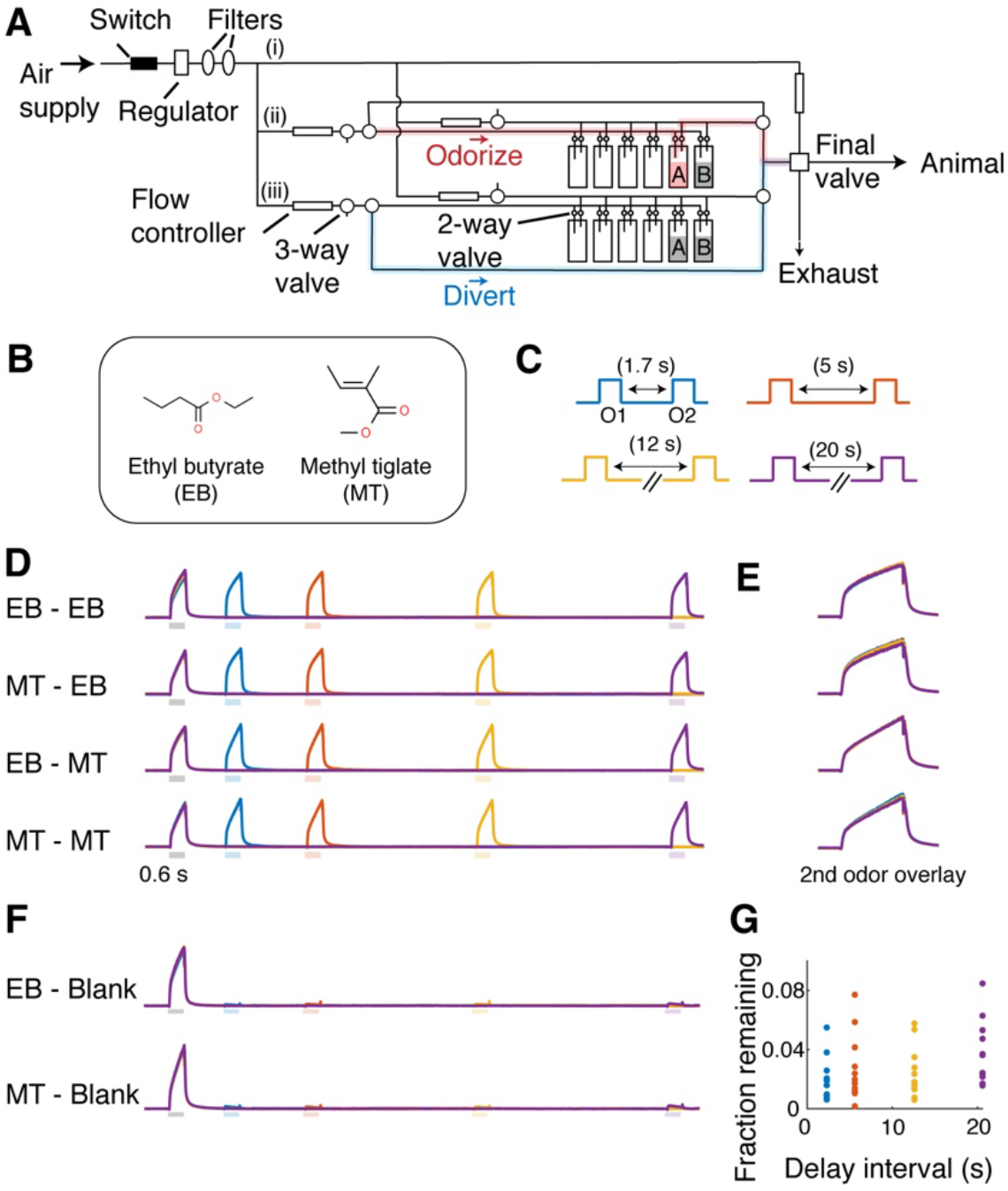
An olfactometer design for stable odour presentations. (**A**) A schematic of the design. Filtered air supply is split into three paths: (i) a stream that normally flows to the animal; (ii) a stream used for the sample odour; (iii) a stream used for the test odour. Each odorising stream has a pair of 3-way valves. When one path is engaged for odour presentation, the air from the selected odour path passes through an odour canister before being directing towards the final valve (red highlight). Simultaneously, the other path diverts air directly towards the final valve (blue highlight), bypassing the odour canisters. (**B**) Odours used in this paradigm. (**C**) Four sample (O1) - test (O2) odour delays used for assessing the olfactometer performance, which were randomly interleaved. (**D**) Example photoionisation (PID) signals for the 4 permutations of sample and test odours (average of 12 trials), color-coded by the interval used as in (**C**). (**E**) PID signals for the test odour overlaid for all intervals tested. (**F**) Test odours were passed through blank odour canisters to test the level of cross-contamination at 4 delay intervals as in (**C**). (**G**) PID signals during the test stimulus periods were expressed as a fraction of the sample odour signal level. Levels from individual trials shown.

The stability of the olfactometer output was tested with a photoionisation detector, which showed that the sample and test odour pulses are stable across various delay durations (Fig. 1B-E). The level of cross-contamination was assessed by measuring the level of residual signals when test odour reservoirs were replaced with empty canisters (Fig. 1F). This indicated minimal cross-contamination from the sample odour (”blank”; Fig. 1F,G; average residual signal as a fraction of the sample odour amplitude = 0.027 ± 0.003; p = 0.17, 1-way ANOVA testing for the effect of delay duration; n = 12 trials per delay). These results indicate that this olfactometer is capable of presenting odours in a manner suitable for the DNMS.

Having designed and tested the olfactometer, we wished to assess if a simple training protocol enables head-fixed mice to acquire a Go/No-Go DNMS paradigm efficiently. Specifically, we tested if auto-association phases, punishments, and multiple training sessions per day used in some implementations could be eliminated. After several days of habituation to head-fixation (median 4 days; n = 6 mice), head-fixed mice generated licks readily to receive the water reward (see Methods). We then subjected the mice immediately to olfactory DNMS training (Fig. 2A). The two odours used in this paradigm were ethyl butyrate (EB) and methyl tiglate (MT). On each trial, the identities of sample and test odours were randomly chosen and presented to the animal with a 5 s delay between odours (Fig. 2). When the sample and test odours did not match (”non-match” trial), a reward (2 droplets of 10 µl water) was delivered unconditionally, with a default delivery at 2.5 s after the test odour offset. However, the reward was delivered earlier if the mice produced early and vigorous anticipatory licks, with the earliest delivery at 0.5 s after the test odour offset. On average, the reward was dispensed when 3 anticipatory licks were detected (see Methods).

**Figure 2:**
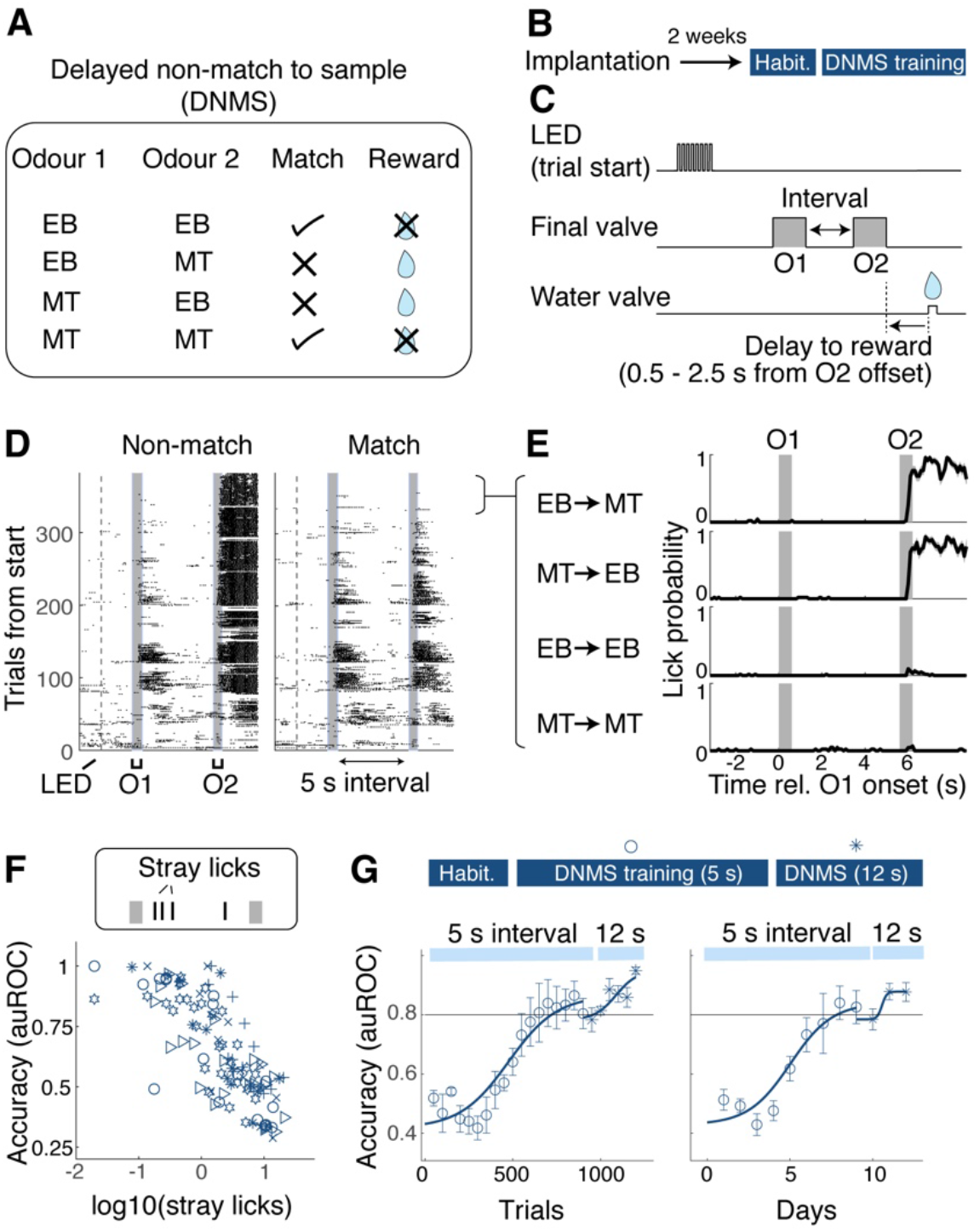
Go/No-Go olfactory delayed non-match to sample with a 5s delay. (A) Illustration of the olfactory delayed non-match to sample (DNMS). On a given trial, two odours (odour1 and odour 2) are presented. Rewarded trials are trials where two odours are not the same (non-match). Reward is 20 ul water. EB = ethyl butyrate, MT = methyl tiglate. (B) Timeline of training. Behavioural training started 2 weeks after surgery for head-plate implantation. (C) Schematic showing a trial structure. Flashes of LED indicate the trial start. Sample and test odours (O1 and O2, respectively) are presented with an interval (5 s for the initial training, and 12 s afterwards). Water reward was delivered on all rewarded trials. Reward was delivered earlier when mice generated anticipatory licks earlier (range of possible reward times = 0.5 - 2.5 s from O2 offset). (D) Lick raster for an example session, separated by non-match trials (left) vs. match trials (right). (E) Example peri-stimulus time histogram of licks separated by 4 trial types from a proficient mouse. (F) Relationship between stray licks (licks during the sample-test odour interval) and accuracy of DNMS performance. Symbols indicate individual mice. (G) Learning curves for 5s delay (hollow) and 12 s delay (stars) shown with respect to the trials from the start of training (block size = 50 trials; left) and with respect to days (right). n = 6 mice. Mean and SEM shown.

In the initial phase of the training, the mice generated many sporadic licks, regardless of the sample and test odour combinations, and also during the sample-test odour delay (Fig. 2D). However, with training, mice learned to produce anticipatory licks more selectively in response to non-matching sample and test odour combinations (Fig. 2E-G). They reached a criterion level of performance (area under Receiver Operator Characteristic (auROC) = 0.8) on average within 705 ± 83.6 trials, or 7.2 ± 0.7 days (mean ± SEM; n = 6 mice). Notably, other undesirable licks, such as licks during the sample-test delay period, decreased in the absence of punishments (average number of licks during 5 s delay = 7.13 ± 1.60 for the first 2 days vs. 0.85 ± 0.39 for the last 2 days of training; p = 0.016, paired t-test, n = 6 mice). In general, the reduction in these sporadic licks and the behavioural accuracy of the task developed together (Fig. 2D-F; Pearson correlation coefficient = -0.785, p = 1.17 × 10^−24^; n = 112 blocks of 50 trials analysed from 6 mice), suggesting that the loss of sporadic licks may occur as a natural consequence of learning the task. Overall, this learning rate is comparable to previous reports of olfactory DNMS paradigms (Han et al., 2018), especially when the shaping phase of those training paradigms is considered. Further, when the sample-test delay period was increased to 12 s, mice performed significantly above chance from the first session (mean accuracy in auROC = 0.848 ± 0.026; p = 3.81 × 10^−5^; n = 6 mice; Student’s t-test for mean auROC = 0.5), demonstrating that the acquired behaviour is generalisable across delay intervals.

The above result indicates that simplified training leads to a robust acquisition of DNMS performance. We next assessed if the sample-test odour delay duration affects the initial acquisition of the task (Fig. 3). The second cohort of mice was trained with the DNMS task with a shorter sample-test odour delay of 1.7 s. All other aspects were held the same as before (Fig. 3A). The mice reached the criterion level of performance, on average, in 449.0 ± 58.3 trials (mean 5 ± 1 days), which is significantly faster than with the 5 s delay (Fig. 3B; p = 0.039, t-test; n = 6 and 5 mice for 5 s delay and 1.7 s delay, respectively). Again, when the sample-test odour interval was increased to 5 s and 12 s, the performance accuracy was significantly above chance from the beginning (average accuracy on first day = 0.87 ± 0.01 auROC for 5 s and 0.81 ± 0.02 for 12 s; p = 1.53 × 10^−5^ and 1.57×10^−4^; Student’s t-test for auROC = 0.5; n = 5 mice). The two cohorts of mice reached the criterion levels of performance at 5 s and 12 s delays subsequently in comparable numbers of trials (Fig. 3C). Overall, these results demonstrate that acquisition of the DNMS task is more efficient with a shorter sample-test interval, but with no advantage when reaching a proficient performance on a longer sample-test delay is required.

**Figure 3:**
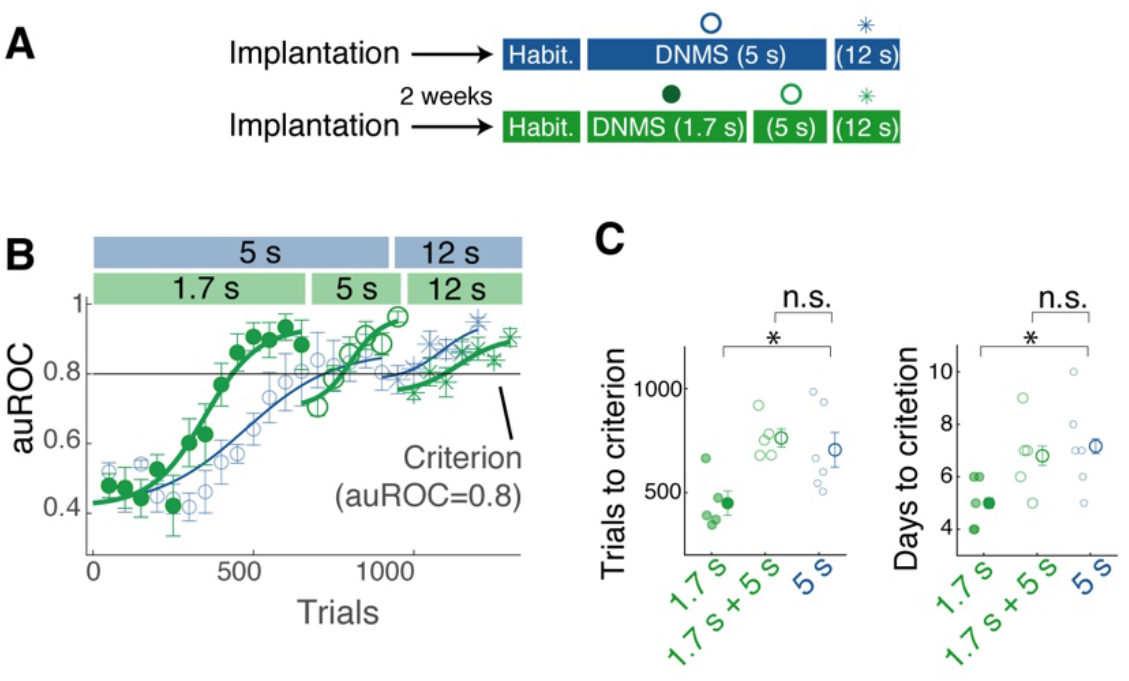
DNMS with a shorter interval is easier to learn. (**A**) Two training designs compared; One cohort of mice that started the training with sample-test odour interval of 1.7 s (green) was compared against those that started with 5 s interval (blue; same data as Fig. 2). (**B**) Learning curves for the two cohorts, showing behavioural accuracy in auROC for for 1.7 s delay (filled circles), 5 s delay (hollow circles) and 12 s delay (stars). Colour scheme as in (**A**). Block size = 50 trials. n = 6 and 5 mice for initial training with 5 s and 1.7 s, respectively. (**C**) Summary comparison of acquisition speeds, in terms of number of trials (left), and number of days (right) taken to reach the criterion (auROC = 0.8). * indicates p = 0.043 (left) and 0.036 (right), 1-way ANOVA with *post-hoc* Tukey-Kramer comparisons. n.s. correspond to p = 0.82 and 0.80 (right). Mean and SEM shown.

In the above training protocols, the reward was delivered earlier when the mice licked earlier to motivate them to perform the task. To determine what influence this reward timing has on the DNMS task acquisition, the third cohort of mice was trained with a fixed reward time while keeping other factors constant, with the initial sample-test delay set to 1.7 s (Fig. 4A,B). The reward (2 × 10 µl water) was delivered 2.5 s after the offset of the test odour regardless of when animals generated anticipatory licks (Fig. 4C,D). On average, the reward was delivered 1.35 ± 0.13 s later for this cohort than for the previous group (p = 6.99 × 10^−7^, 1-way ANOVA; n = 5 and 6 mice for the conditional and fixed reward time groups, respectively). Surprisingly, the learning curves of the two cohorts revealed that the mice that received the reward at a fixed time acquired the task faster than the cohort with conditional timing (Fig. 4E,F; trials to criterion 305.8 ± 30.1 trials, compared to 449.0 ± 58.3 trials for the group with conditional reward timing). However, their ability to perform longer sample-test odour intervals was comparable (average accuracy for the initial sessions = 0.87 ± 0.02 and 0.87 ± 0.03 for 5 s and 12 s intervals, respectively; p = 0.22 and 0.19, 2-way ANOVA for the effect of cohort and interval durations, respectively). Thus, the reward timing affects the task acquisition but not the generalisability of performance across intervals, revealing an unexpected and detrimental effect of early reward delivery.

**Figure 4:**
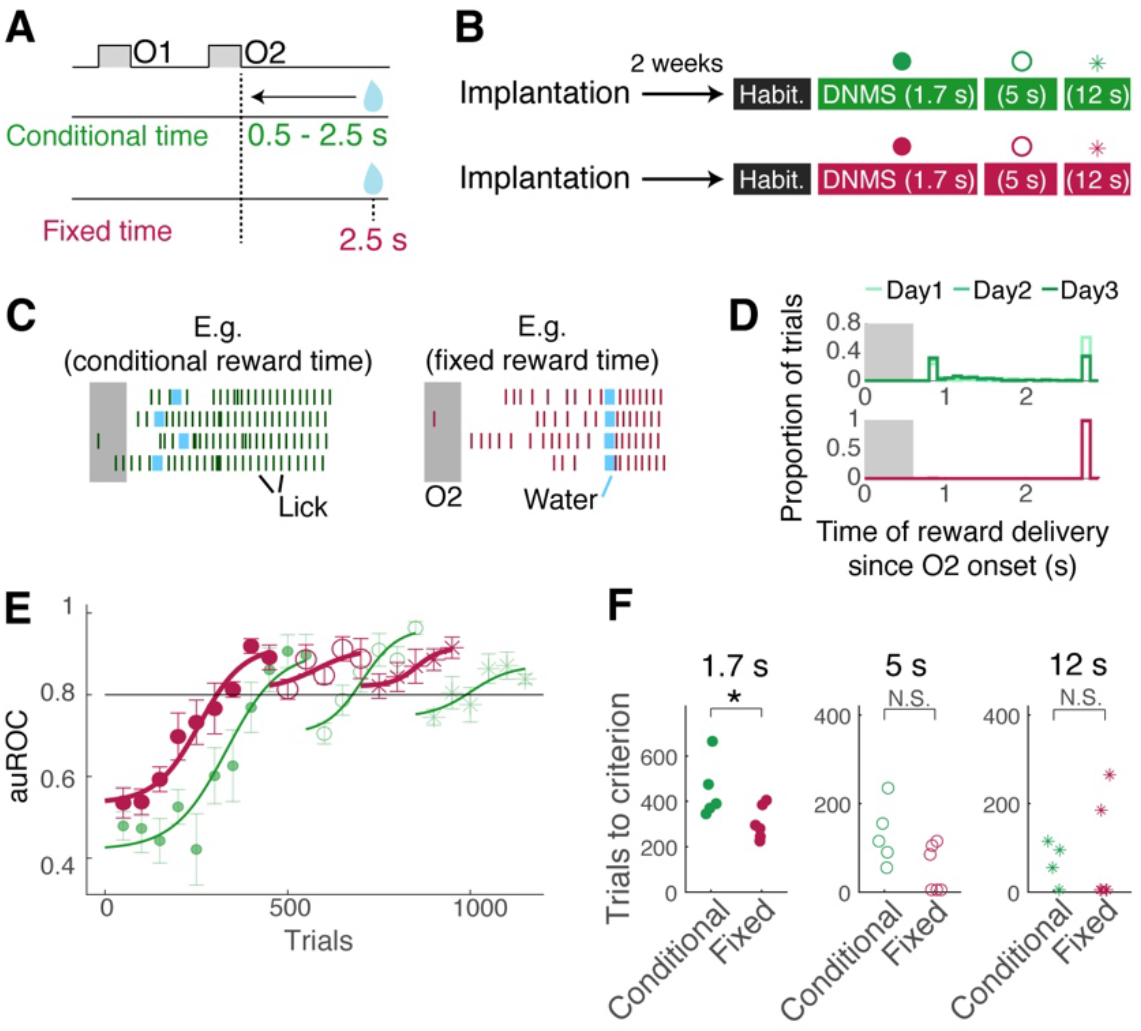
Effect of reward timing on DNMS acquisition. (**A**) With conditional reward timing (green), the water reward was delivered earlier if mice licked earlier (possible reward onset ranged 0.5 - 2.5 s after the odour 2 offset). With fixed reward delivery time (burgundy), water reward was delivered at 2.5 s after the odour 2 offset. (**B**) Timeline of training for the two cohorts, with sample-test odour interval indicated inside the brackets. (**C**) Examples of lick timing (vertical lines) relative to odour 2 (duration 0.6s) and water delivery (light blue). For the conditional reward time group (left), the timing of reward onset depended on lick timing. For the fixed reward onset (right), water timing was fixed regardless of licking behaviour. (**D**) Histograms of water onset times for the two groups, for days 1-3 of the training. (**E**) DNMS learning curves for the two cohorts for 1.7 s (filled circles), 5 s (hollow circles) and 12 s (stars) intervals. Horizontal line indicates the criterion level (auROC = 0.8). (**F**) Summary of learning speeds, measured by number of trials taken to reach criterion performance at sample-test odour interval of 1.7 s (left), 5 s (middle), and 12 s (right) for the two cohorts. p = 0.047, 0.068, 0.87 (two-sample t-tests), respectively. n = 5 mice for conditional reward timing and 6 mice for fixed reward timing.

How do the different protocols affect the quality of the task performance? It is known that a hallmark of working memory is that it lasts briefly and degrades over time (Baddeley, 2011; Blough, 1959; Liu et al., 2014). To test if different protocols lead to DNMS task performance with different characteristics, we assessed the ability of trained mice to generalise the task performance over different sample-test odour intervals within a given session. This included an inter-odour delay interval that the animals had not encountered before. For each trial, the sample-test odour delay duration was randomly chosen from 4 different intervals (1.7 s, 5 s, 12 s, and 20 s; Fig. 5). An analysis of the behavioural performance indicated that the performance was indeed poorer at longer intervals (Fig. 5D). However, no statistically significant difference was present across two cohorts that underwent different training paradigms (p = 2.55 × 10^−6^ and 0.24, 2-way ANOVA for the effects of interval and training history, respectively; n = 4 mice for both cohorts). Overall, the result indicates a greater difficulty with longer intervals but with similar properties, irrespective of how the mice acquired the task.

**Figure 5:**
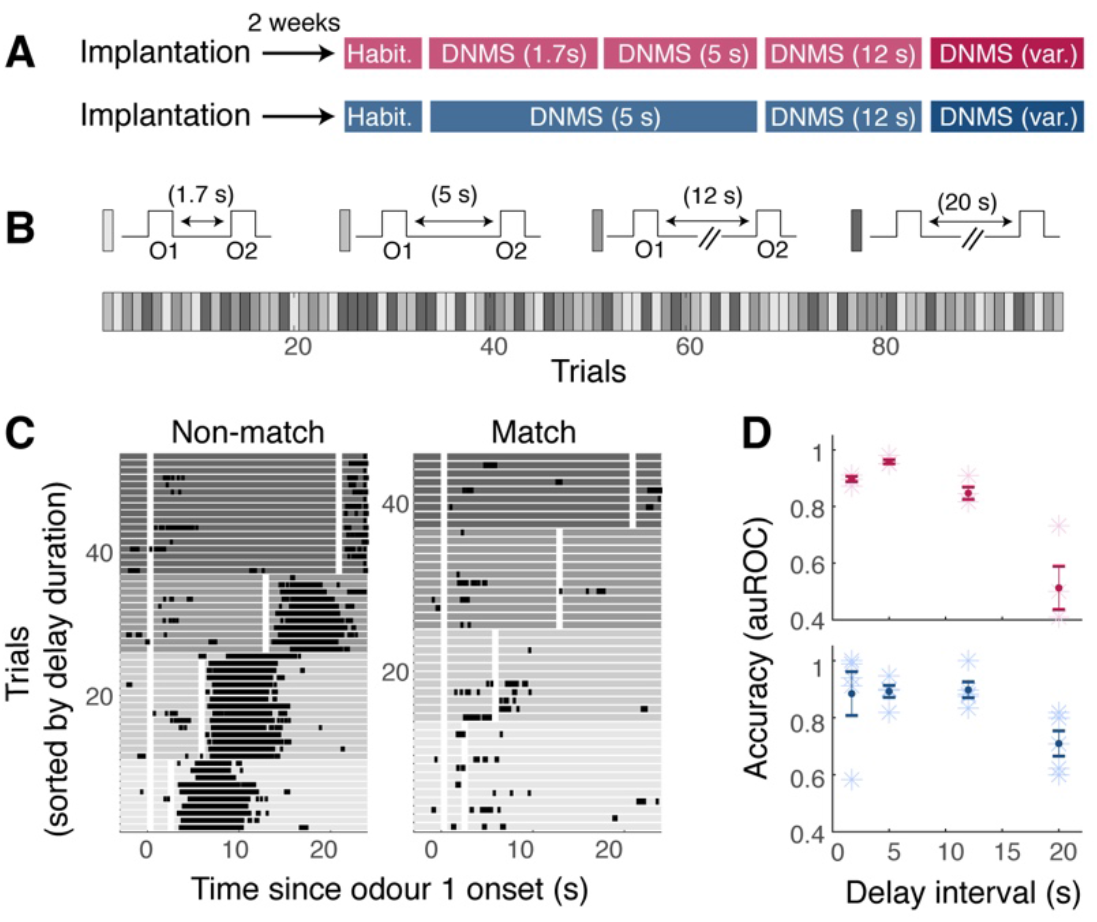
Interval-dependent performance and generalisation. (**A**) Once mice were trained to perform DMNS with a 12 s interval, they went through one session where the sample-test odour interval was randomly selected from 1.7 s, 5 s, 12 s, or 20 s on a given trial (”DNMS (var.)”). (**B**) Example trial order, where grey intensity indicates the sample-test odour interval duration. (**C**) Lick raster, sorted by non-match (left) and match (right) trials, and by the duration of sample-test odour intervals, with grey scale indicating the interval duration. White shaded areas indicate the time of odour presentations. (**D**) Accuracy for each interval used, for two cohorts, with colours corresponding to the cohorts described in (**A**). Mean and SEM shown. n = 4 mice for each cohort.

Having verified that the quality of acquired behaviour is similar across the training methods, we sought to determine why the reward timing affected the initial acquisition rates. Specifically, we wished to understand why the late reward arrival resulted in faster acquisition of the DNMS task. To obtain clues, we first analysed if there was any difference in the stray lick patterns across the three cohorts during learning. We measured the stray licks generated during the sample-test odour delay (Fig. 6A). Among the two cohorts of mice with conditional reward timing, there tended to be more sporadic licks during the sample-test odour interval during acquisition (average number of stray licks per second in the first 400 trials = 1.17 ± 0.29, 1.58 ± 0.21, and 0.61 ± 0.14 for cohorts 1, 2, and 3, respectively; p = 0.018; 1-way ANOVA; n = 6,5, and 6 mice, respectively). Second, we noticed that the reaction times - i.e., how quickly the mice generated anticipatory licks in response to the test odour presentation - differed between cohorts with reward timing differences. The two cohorts that could receive reward early gradually shifted to shorter reaction times with training, while the cohort with fixed and later reward timing tended to maintain late lick onsets (Fig. 6A,C,D; average reaction times for the last 100 reward trials = 0.65 ± 0.07 s, 0.78 ± 0.12, and 1.78 ± 0.11 s for two cohorts with conditional reward timing, and the cohort with fixed reward timing, respectively; p = 1.43 × 10^−6^, 1-way ANOVA). Third, when the false alarm patterns were analysed, we found more erroneous licks in the cohorts with early reward (Fig. 6E). Altogether, these traits indicate that the conditional reward timing makes mice generate behavioural outputs more readily, indicating a lowering of the internal threshold for generating licks.

**Figure 6:**
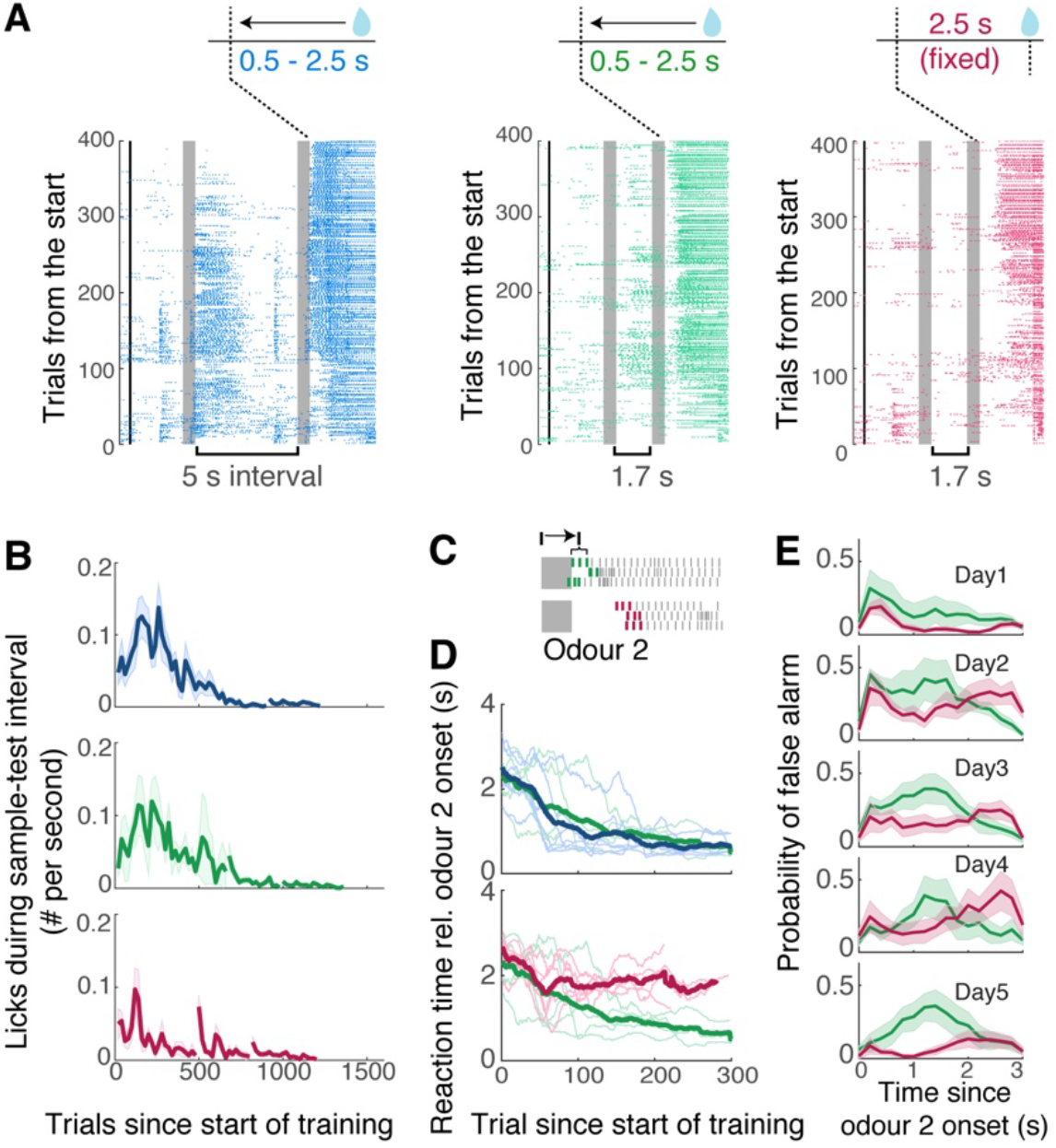
Behavioural output patterns depend on reward timing. (**A**) Example lick raster plots from mice that went through different training paradigms: left, a mouse that started with 5 s sample-test odour interval and conditional reward timing; middle, starting with a 1.7 s interval and conditional reward timing; right, starting with a 1.7 s interval and fixed reward timing. (**B**) Evolution of stray licking during sample-test odour interval for the three cohorts (n = 6, 5, and 6 mice, respectively). Thick lines = mean; shades = mean ± SEM. (**C**) Illustration of a reaction times: timings of first 3 licks after the odour 2 onset are averaged and expressed relative to the onset of odour 2. (**D**) Evolution of reaction time with training for the three cohorts indicated with the colour scheme as in (**A**). Thin lines = individual mice, thick lines = average. (**E**) Peri-stimulus time histograms for licks generated on match (unrewarded) trials were compared between cohorts that started with the 1.7 s delay but differed in the reward timing.

To analyse the relationship between the reward timing and internal threshold more rigorously and to explain how a lower behavioural threshold relates to slower learning, we analysed our data using a simple drift-diffusion model (Ratcliff and McKoon, 2008). This model simulates the internal representation of sensory evidence as an accumulation of noisy diffusion processes (”drift rate” with noise; Fig. 7A). In addition, the time at which this sensory evidence crosses a threshold (”bound”) models the behavioural reaction times observed. Here, we fit animals’ reaction times and estimated two diffusion rates, corresponding to the strengths of momentary sensory drives for matching and non-matching odour stimuli (S- and S+ stimuli, respectively), as well as a single bound (”Go bound”) (Ratcliff et al., 2018). The fitted model captured the tendency for early reaction times, as well as more frequent false alarm occurrences, for the cohort with conditional reward timing and more accurate and later production of anticipatory licks for the cohort with fixed reward timing (Fig. 7B).

**Figure 7:**
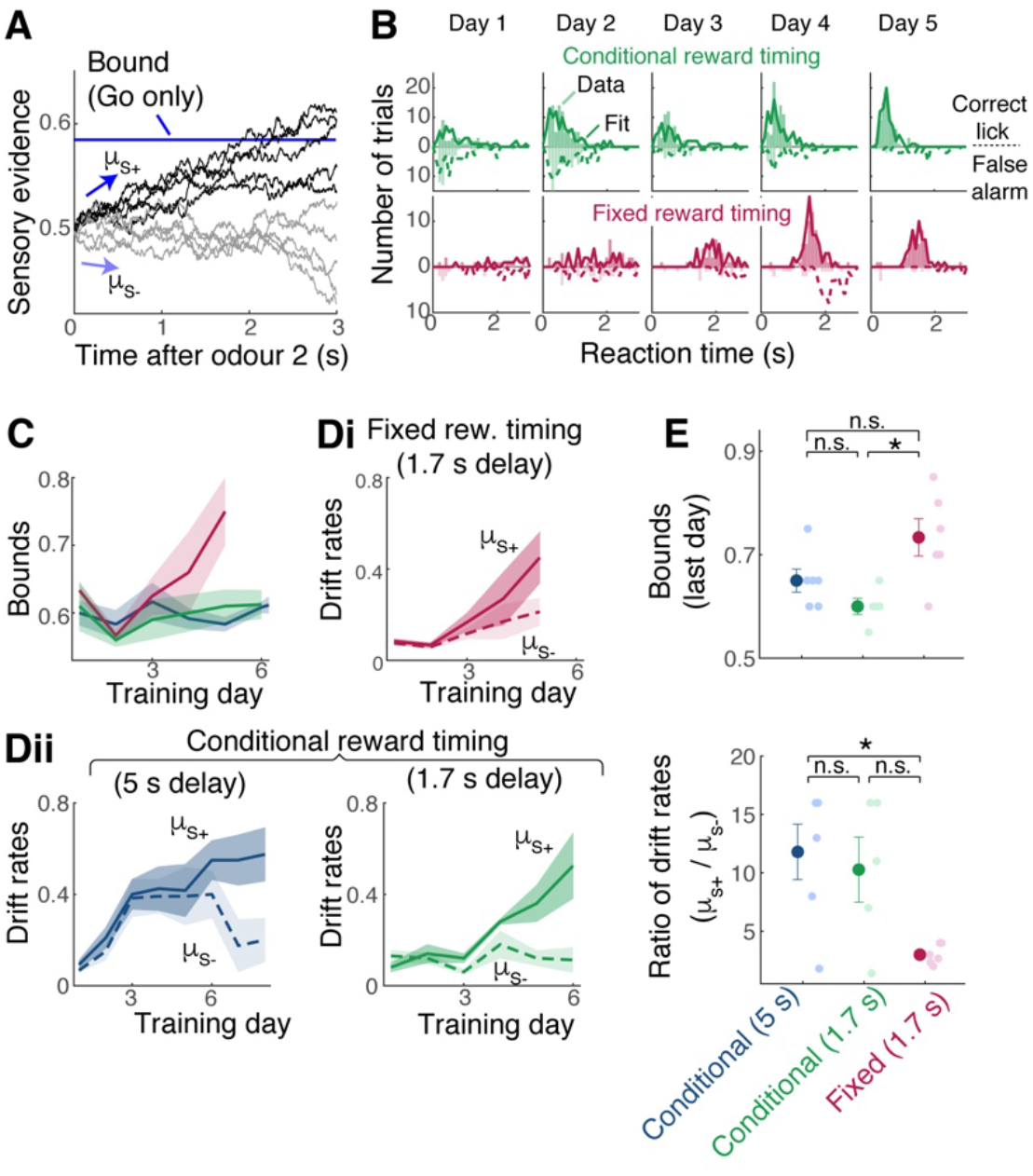
Interplay between reward timing, behavioural threshold, and the discriminability of sensory representations. (**A**) A drift diffusion model with one bound (blue line) to model the Go/No-Go behaviour. Drift rates, µ_s+_ and µ_s-_, are the strengths of momentary evidence following the S+ (non-match) stimulus and S-(match) stimulus, respectively. At each time point, fixed amount of noise is added, and accumulated over time (sensory evidence). Reaction time is the time at which the sensory evidence crosses the bound. (**B**) Histograms of reaction times for an example animal over 5 training days. Observed licks (correct licks = licks on S+ trials and false alarms =licks on S-trials) shown as bars. Simulated result using fitted parameters superimposed with lines (solid lines = S+ simulation, dotted lines = S-simulation). (**C**) Estimated bounds for cohorts with fixed reward timing (burgundy) and conditional reward timing (green). (**Di**) Estimated drift rates for the cohort with late, fixed reward timing. (**Dii**) Estimated drift rates for two cohorts with conditional reward timing. (**E**) Comparison of estimated bounds (top) and the ratio of drift rates (µ_s+_/µ_s-_; bottom) for the three cohorts, from the last day of initial training. Statistics: for decision bounds, p = 0.011 and 0.099 across reward timing and 0.43 for cohorts with same reward timing but different sample-test intervals (one-way ANOVA with *post-hoc* Tukey-Kramer tests); for drift rates, p = 0.0183 and 0.0639 across reward timing, and 0.86 for cohorts with same reward timing (one-way ANOVA with *post-hoc* Tukey-Kramer tests). Mean and SEM shown.

This analysis revealed two main ways the reward timing may affect the acquisition of a task. First, as indicated by the readiness to generate early anticipatory licks and sporadic licks, earlier arrival of reward tends to lower the decision bound (Fig. 7C; Mean bounds = 0.65 ± 0.02 and 0.60 ± 0.02 for the two cohorts with early reward vs. 0.73 ± 0.04 for the fixed, late reward group; p = 0.013, 1-way ANOVA; n = 5, 6, and 6 mice). Second, we found that the difference in the drift rates associated with the S+ and S-stimuli became greater for the cohort with conditional reward timing (Fig. 7D,E; ratio of S+ vs. S-diffusion rates = 11.8 ± 2.37 and 10.28 ± 2.8 for the two cohorts with conditional reward timing vs. 3.00 ± 0.34 for the fixed, later reward group; p = 0.017, 1-way ANOVA; n = 6, 5, and 6 mice). In other words, when the reward arrives early, a consequence is a lowering of the decision threshold, which may require the sensory representations to be more discriminable to achieve the same accuracy. This requirement for more discriminable representation may underlie the slower task acquisition.

## Discussion

Describing the quantitative relationships between physical stimuli and behavioural change is central to gaining insights into learning. In this study, we demonstrate that simplified training, without punishment, leads to a robust acquisition of the olfactory DNMS task. Further, we identified two aspects of stimulus timing that affect the task acquisition rate. The task acquisition is faster when the sample-test delay interval is shorter and slower when animals could receive a reward earlier.

The effect of the delay interval on the DNMS task acquisition may relate to the degradation or fading of the short-term memory over time, which is a hallmark of working memory (Baddeley, 2011). An underlying neural correlate of short-term information retention is thought to be a stimulus-specific, persistent increase in firing rates in individual neurons. Such activity patterns are prevalent in the prefrontal cortex in animals engaged in delayed response and delayed matching tasks (Funahashi et al., 1989; Fuster and Alexander, 1971; Goldman-Rakic, 1995; Miller et al., 1996; Wu et al., 2020). These activities may be maintained through dynamic interactions within a local network (Goldman-Rakic, 1995; Sreenivasan and D’Esposito, 2019) and possibly across brain regions (Wu et al., 2020; Zhang et al., 2019). However, typically, there is a degradation - or decay in the firing rate - during the delay period, which correlates with task performance (Zhang et al., 2019). Thus, the faded representation may slow down the task acquisition in a manner similar to presenting a weaker stimulus in associative learning (Grant and Schneider, 1948). It should be noted that, while this study used a Go/No-Go paradigm, such a weakening of sensory representations is likely to have a similar effect on other instances of delayed (non-)match-to-sample, such as the two-alternative forced choice paradigm.

The detrimental effect of early reward arrival on task acquisition was less expected from the outset. For example, in an influential framework, the temporal contiguity between stimuli and reinforcing signals is described by an eligibility trace, a short-term memory vector that signals a magnitude of reinforcement learning permitted (Barto et al., 1983; Klopf, 1972). Since this describes a signal that fades over time, a simple prediction is a slower learning rate with later reward arrival. Further, some studies suggest that waiting longer or more prolonged evidence accumulation can concur a cost to behavioural accuracy (Drugowitsch et al., 2012). Indeed, when the reward could be delivered earlier, our mice also developed early licking. This effect on reaction times was interpreted as lowering the decision bound. A consequence of this lower bound seems that to achieve the same accuracy, the momentary strengths of the sensory drive (the “drift rates”) had to become more divergent. This requirement for more divergent sensory drives may underlie a slower task acquisition.

While the drift-diffusion model was instrumental in narrowing potential factors involved, what might the abstract, computational terms correspond to physiologically? Several studies have reported ramping neuronal activities in prefrontal and frontal cortices, for example, in monkeys discriminating and reporting the direction of random dot motions with saccades after delays (Ding and Gold, 2012; Roitman and Shadlen, 2002; Shadlen and Newsome, 1996). Furthermore, the slopes of such ramping activity are known to increase with the coherence of motion or the strength of stimuli (Ding and Gold, 2012; Roitman and Shadlen, 2002; Shadlen and Newsome, 1996). These suggest that the drift rates of the model may be interpreted in terms of mechanisms that lead to action potential generation in relevant prefrontal cortex neurons, for example, the firing rates of neurons presynaptic to these neurons.

In the case of the olfactory delayed matching-to-sample, a recent study suggested that the anterolateral motor (ALM) cortex may contain neurons that report the match/non-match of sample and test odours (Wu et al., 2020). According to their model, the persistent, stimulus-specific activity in response to the sample odour is maintained through recurrent interactions involving many sensory cortices, which is used to compare against the test odour identity to compute the match in ALM. We, therefore, speculate that the drift rate may correspond to this process. Furthermore, with learning, the selectivity of choice-related activity may undergo refinement, just as sensory representations in many brain regions are known to increase in selectivity with the acquisition of a task (Poort et al., 2015).

In summary, our results indicate two ways in which stimulus timing affects stimulus encoding in relation to the acquisition of a DNMS task: retention of sensory information over time and discriminability of match-related signals required for accurate behavioural performance. The efficient DNMS training paradigm we describe here, in turn, may accelerate the investigations of underlying neural mechanisms.

## Methods

### Animals

All animal experiments had been approved by the OIST Graduate University’s Animal Care and Use Committee (Protocol 2021-350). C57Bl6J mice were purchased from Japan CLEA (Shizuoka, Japan) and were acclimatised to the facility for at least 1 week before they were used for experiments. All mice used in this study were adult males (8 – 11 weeks old at the time of head plate implantation). Littermates were used in the three cohorts. Mice were randomly distributed across cohorts and all cohorts were trained on the same days, with staggered sessions per day.

### Olfactometry

A custom-made flow-dilution olfactometer was used to present odours. Briefly, custom Labview codes were used to control: (1) sourcing output digital output modules (NI-9474, National Instruments), which actuated direct-operated solenoid valves. These were used to control air flow and gate reward delivery (water droplets) delivery; (2) analog output modules (NI-9263, National Instruments) to generate signals for flow controllers (C1005-4S2-2L-N2, FCON, Japan) and to communicate the output trial types with acquisition boards; and (3) digital input/output modules (NI-9402, National Instruments) to control LEDs and to communicate valve opening timing.

A pair of normally closed solenoid valves was assigned per odorant to odorise the air stream. These solenoid valves were attached to a manifold, such that a set of 8 pairs connected to the common air stream (Fig. 1). On each trial, the sample and test odors came from two separate manifolds to avoid cross-contamination. Further, each odorant was duplicated on two manifolds to allow presentation of a particular odour from either manifold, avoiding an auditory association for the behavioural task. An air stream was odorised only when a pair of three-way valves were actuated and directed towards the animal when the solenoid valve closest to the animal (final valve) opened. The final valve was opened for 0.6 s for each odour presentation. Odours were presented at approximately 1 % of the saturated vapor. Total air flow, which is a sum of odorised air and the dilution air, was approximately 2 L/min, which was matched by the air that normally flows towards the animal. The inter-trial interval was approximately 40 seconds in order to purge the airways to minimise cross-contamination. Ethyl butyrate (W242705) was purchased from Sigma-Aldrich and methyl tiglate (T0248) was from Tokyo Chemical Industry (Tokyo, Japan). Stock odorants were stored at room temperature in a cabinet filled with N_2_ and away from light.

### Surgery

Head plate implantation: All recovery surgery was conducted in an aseptic condition. 8 to 11-week-old male C57Bl6/J mice were deeply anaesthetised with isoflurane. The body temperature was kept at 36.5 °C using a heating blanket with a DC controller (FHC, Bowdoin, USA). To attach a custom head plate about 1 cm in width weighing a few grams, the skin over the parietal bones was excised, and the soft tissue underneath was cleaned, exposing the skull. The exposed skull was gently scarred with a dental drill, cleaned, dried, and coated with cyanoacrylate (Histoacryl, B.Braun, Hessen, Germany) before placing the head plate, which was fixed in place with dental cement (Kulzer, Hanau, Germany). Mice were recovered in a warm chamber, returned to their cages, and given carprofen subcutaneously (5 mg/kg) for 3 consecutive days.

### Habituation and behavioural measurements

Water restriction began 2 to 3 weeks after surgery. Mice went through habituation to head fixation, one session per day for approximately 30 minutes. No odours were presented. This lasted 3 days or until the mice learned to lick vigorously for a total water reward of at least 1 ml. The respiration pattern was measured by sensing the airflow just outside the right nostril by placing a flow sensor (AWM3100V, Honeywell, North Carolina, USA). Lick responses were measured using an IR beam sensor (PM-F25, Panasonic, Osaka, Japan) that was a part of the water port. Nasal flow, an analogue signal indicating the odours used, lick signal, a copy of the final valve and water valve timing were acquired using a multifunction I/O device (USB-6363; National Instruments).

### Olfactory DNMS training

After habituation, the head-fixed mice were trained to associate a water reward with a target odour (ethyl butyrate). The reward comprised two droplets of water (10 µl each), that arrived after the onset of the final valve opening. Correct reactions were defined as the generation of anticipatory licks in response to the test odour that did not match the sample odour, and a lack of licking in response to the test odour when it is the same as the sample odour. Once the overall accuracy was above 80% in at least one behavioural session, the delay interval was increased. A typical training session comprised roughly 100 trials, lasting about 1 hour. Rewarded and unrewarded trials occurred at equal probability, where the specific permutation of the odours used (e.g., EB-MT or MT-EB for rewarded trials) was randomly assigned. To deliver reward with a timing that depended on the timing of mouse’s anticipatory licks, lick signals were analysed online for a brief period following the test odour presentation (0.6 - 3 s after the test odour onset). The reward was delivered on rewarded trials when the number of anticipatory licks exceeded a threshold. The threshold for triggering an output here was equivalent to 3 licks. Inter-trial interval (interval between the test odour onset and the start of the next LED signal) was approximately 18 s. All data were acquired at 1 kHz.

### Data analysis

#### Analysis of the behavioural data

The behavioural data were analysed using built-in event detection functions in Spike2 to obtain valve opening times and further analysed in Matlab using custom codes. The number of anticipatory licks after the test odour offset was counted for each trial to calculate the accuracy. When the reward timing was variable, the temporal window analysed was until the onset of the water valve opening. To analyse comparable temporal windows across trial types, for unrewarded trials, an average water valve timing from the rewarded trials was used. For fixed reward timing, a 2.6 s window from the test odor offset was used. Using a shorter window to mimic the condition for the variable reward timing did not affect the result. To calculate the learning curve, the accuracy was expressed as the area under the Receiver Operator Characteristic (auROC) in a given block of 50 trials, using the Matlab function *perfcurve*.

##### Trials to criterion

For each animal, block-by-block auROC values were fitted with a logistic function. Using this fit, the number of trials required to reach auROC value of 0.8 was interpolated for each animal.

##### Reaction time

The reaction time was calculated by measuring the timings of the first 3 licks after the test odour on rewarded trials, expressed relative to the onset of the test odour, and averaged.

##### Stray licks

All licks between the onsets of the sample and test odours were counted and divided by this interval. Probability of false alarms: lick occurrences following the test odour presentation (0 - 3 s relative to the onset) on unrewarded trials were detected and used to construct peri-stimulus time histograms with a bin size of 0.02 s, which was normalised by the number of trials.

#### Drift diffusion model

The reaction times were modelled using a simple drift-diffusion model is based on (Ratcliff and McKoon, 2008) but adapted for the Go/No-Go paradigm as described in (Ratcliff et al., 2018). The model is described by three parameters: the Go decision bound and two instantaneous drift rates, ε ξπρεσσεδ ασ µ_s-_ for match (S-) and µ_s+_ for non-match (S+) trials. At time t = 0, the accumulated evidence was set to 0.5 for all data. Time steps were discretised in 20 ms (Δt). For each new time step, new momentary evidence was drawn from a stationary distribution: x(Δt) = N(µ_i_,1), a normally distributed random generator with a mean of µ_i_ where i = [S+, S-], and a noise parameter of 1 standard deviation. This was accumulated over time to yield the sensory evidence: 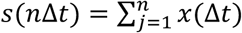. The reaction time was defined as the time at which this sensory evidence crossed the decision bound. The parameters were obtained for each animal by fitting the data from each behavioural session by minimising the chi-square values (Ratcliff et al., 2018).

## Acknowledgement

We thank Yu-Pei Huang and OIST’s Animal Resource Service staff for their dedicated technical assistance, members of the Sensory and Behavioural Neuroscience Unit for discussions, and Sander Lindeman for comments on the manuscript. We are grateful for the support from the OIST Graduate University.

## Author contributions

Conceptualisation, JR, JKR and IF; Investigation, JR and JKR; Data curation, JR and JKR; Analysis, JR, JKR, and IF; Writing, JR, JKR, and IF.

